# Multi-site internal modification of long DNA substrates for single-molecule studies

**DOI:** 10.1101/045955

**Authors:** Armando de la Torre, Yoori Kim, Andrew A. Leal, Ilya J. Finkelstein

## Abstract

Single-molecule studies of protein-nucleic acid interactions frequently require site-specific modification of long DNA substrates. DNA isolated from bacteriophage λ (λ-DNA) is a convenient source of high quality long (48.5 kb) DNA. However, introducing specific DNA sequences, tertiary structures, and chemical modifications into λ-DNA remains technically challenging. Most current approaches rely on multi-step ligations with low yields and incomplete products. Here, we describe a molecular toolkit for rapid preparation of modified λ-DNA. A set of PCR cassettes facilitates the introduction of recombinant DNA sequences into λ-DNA with 90-100% yield. Furthermore, various DNA structures and chemical modifications can be inserted at user-defined sites via an improved nicking enzyme-based strategy. As a proof-of-principle, we explore the interactions of Proliferating Cell Nuclear Antigen (PCNA) with modified DNA sequences and structures incorporated within λ-DNA. Our results demonstrate that PCNA can load on both 5’-ssDNA flaps and a 13xCAG triplet repeat. However, PCNA remains trapped on the 13xCAG structure, confirming a proposed mechanism for triplet repeat expansion. Although we focus on λ-DNA, this method is applicable to all long DNA substrates. We anticipate that this molecular toolbox will be broadly useful for both ensemble‐ and single-molecule studies that require site-specific modification of long DNA substrates.

## Introduction

Single-molecule imaging and manipulation approaches have greatly expanded our understanding of protein-nucleic acid interactions (1). More recently, the single-molecule toolkit has expanded to include hybrid methods that combine fluorescence imaging (e.g., single-molecule FRET) with force manipulation (e.g., optical or magnetic tweezers) for multi-modal data acquisition (2–4). Many of these approaches require very long (>10 kb), chemically modified DNA substrates. Long DNA substrates are frequently used as flexible, micron-long tethers in optical and magnetic tweezers experiments (4). These substrates are especially useful for observing proteins that traverse along DNA and have been used for characterizing enzymes that package, transcribe, replicate, or diffuse on DNA (5). Moreover, long DNA molecules elevate the biochemical reactions away from glass coverslips and other surfaces, reducing the potential for non-specific interactions (6). DNA derived from bacteriophage λ (λ-DNA) offers several advantages for single-molecule studies: it is a long (48.5 kb) substrate, permitting the observation of protein movement over kilobase-length distances. In addition, the *cosL* and *cosR* ssDNA overhangs facilitate direct ligation of modified DNA handles. Finally, high-quality recombinant DNA can be purified from lysogenic cells in large quantities.

Site-specific λ-DNA modification strategies generally fall within one of four categories: (i) restriction enzyme cleavage and ligation (4, 7); (ii) recombinase-mediated modifications (8); and (iii) insertion of extrahelical structures via oligonucleotide mimics, or (iv) nicking endonuclease (nickase)-based oligo replacement (9–13). To date, restriction enzyme cleavage and multi-step ligation *in vitro* is one of the most frequently used methods for modifying λ-DNA. However, this approach is technically challenging because multi-step intra-molecular ligation is inefficient. Restriction enzyme-based cloning is also limited due to the few unique restriction sites within the 48.5 kb phage genome. Furthermore, producing appreciable quantities of recombinant DNA requires packaging the λ-DNA into phage particles, several rounds of viral amplification, and infection on an *E. coli* host with high-titer λ-phage prior to DNA purification.

Site-specific recombination *in vitro* is another promising approach for inserting exogenous DNA structures into λ-DNA (8). However, this approach requires commercially unavailable enzymes. In addition, two unique recombinase sites must also be cloned into λ-DNA at the desired insertion position. Introducing site-specific synthetic oligonucleotides and extrahelical structures poses additional challenges in such a long DNA substrate. Several groups have used locked‐ and peptide-nucleic acid oligonucleotides (LNAs and PNAs, respectively) to label specific sites within λ-DNA via formation of triplex-forming DNA motifs (11–14). However, these methods rely on non-physiological structures, require careful selection of the appropriate hybridization site, and need multiple rounds of optimization with expensive non-canonical oligonucleotides. Finally, a nicking endonuclease (nickase) based strategy has been reported for inserting synthetic oligonucleotides into both plasmids and λ-DNA (10, 15–17). Nickase-based insertion is a versatile tool for modifying plasmid-length DNA (18), but is less appropriate for λ-DNA because of the large number of nicks that most commercially-available enzymes leave in the DNA substrate.

Here, we develop a method for more rapidly modifying and purifying recombinant λ-phage DNA. We use *in vivo* recombineering to target any segment of the phage genome, abrogating the need for restriction sites and ligation. Using this approach, we develop a rapid molecular toolkit for inserting exogenous DNA sequences at three unique sites with 90-100% efficiency. Site-specific DNA incorporation is confirmed via both ensemble and single-molecule fluorescence assays. We also demonstrate a strategy for inserting non-replicative extrahelical DNA structures at these sites. We explore the utility of these DNA structures by demonstrating site-specific loading of PCNA by the clamp-loader complex Replication Factor C (RFC). Our results show that PCNA can be loaded on both flaps and a 13xCAG triplet nucleotide repeat (TNR). Surprisingly, PCNA remains trapped within the 13xCAG repeat, adding further support to a model that suggests PCNA participates in TNR expansion via illegitimate activation of the DNA mismatch repair machinery (19, 20). In sum, we anticipate that this molecular toolkit will be broadly useful for both ensemble and single-molecule studies that require site-specific modification of long DNA substrates.

## MATERIALS AND METHODS

### Proteins and DNA

FLAG-epitope labeled Lac repressor was purified from BL21(DE3) as described previously (21). Biotinylated anti-FLAG M2 antibodies were purchased from Sigma and streptavidin-conjugated quantum dots (QDs) were purchased from Life Technologies. All oligonucleotides were ordered form IDT and are summarized in Table S1. Plasmids containing an antibiotic cassette flanked by two 200-basepair segments that are homologous to the λ-phage genome were custom synthesized by BioMatik. All plasmid modifications were carried out using the Q5 inverse PCR mutagenesis kit (NEB).

### Replication Factor C (RFC)

The plasmids pLant2b-RFC-AE and pET11-RFC-BCD were kindly provided by Manju Hingorani (22). The two plasmids were co-transformed into BL21(DE3) ArcticExpress cells (Agilent). A single colony was inoculated into 680 μL of LB and grown overnight at 37°C. 200 μL of these cells were then seeded into 2 L LB with 50 μg mL^−1^ kanamycin and 50 μg mL^−1^ carbenicillin. The culture was grown at 30°C to an OD_600_~0.6, cooled on ice with swirling to 16°C and induced with 0.5 mM IPTG. Induction continued for 16 hours at 12°C. Cells were harvested by centrifuging for 15 minutes (3,300 RCF at 4°C) and resuspended in 20 mL of lysis buffer (30 mM HEPES [pH 7.5], 0.25 mM EDTA, 5% (v/v) glycerol). The resuspended cell paste were either frozen in liquid nitrogen and stored at −80°C, or prepared for lysis by adding 250 mM NaCl, 1 mM phenylmethanesulfonyl fluoride (PMSF, Sigma-Aldrich), and 1x HALT protease inhibitor (Thermo-Fisher). Cells were lysed in a homogenizer (Avestin), and centrifuged (140,000 RCF, at 4°C) for 35 minutes. A home-made 7 mL SP FF column (resin from GE Healthcare) was equilibrated with 35 mL of buffer A (30 mM HEPES [pH 7.5], 0.25 mM EDTA, 5% (v/v) glycerol) + 200 mM NaCl and the clarified lysate was loaded onto the equilibrated column at 0.5 mL min^−1^. The column was then washed with 35 mL of buffer A + 250 mM NaCl and the protein was eluted with a 100 mL gradient to buffer B (30 mM HEPES [pH 7.5], 0.25 mM EDTA, 5% (v/v) glycerol, 1 M NaCl) at 1 mL min^−1^. RFC-containing fractions were pooled and developed through a 1 mL Q HP column (GE Healthcare) preequilibrated with buffer A + 100 mM NaCl. The combined sample from the SP column was loaded onto the Q column at a rate of 0.7 mL min^−1^. The column was washed with 10 mL of buffer A + 110 mM NaCl at 0.8 mL min^−1^, and then eluted with an 18 mL gradient into buffer B at 0.8 mL min^−1^. The eluted Q column fractions were analyzed on a 10% SDS-PAGE gel, and the fractions containing RFC were combined, frozen in liquid nitrogen, and stored at −80°C. The RFC concentration was determined by comparison to a BSA titration curve using SDS-PAGE.

### Proliferating Cell Nuclear Antigen (PCNA)

A plasmid expressing *Saccharomyces cerevisiae* His6-PCNA was kindly provided by Francisco Blanco (23). A triple FLAG epitope was introduced at the N-terminus via inverse PCR mutagenesis (NEB) using primers YK_PCNA01 and YK_PCNA02 (see Table S1) to generate pIF105.pIF105 was transformed into BL21(DE3) codon plus RIL cells. A colony was inoculated into 30 mL LB with 50 μg mL^−1^ kanamycin and 34 μg mL^−1^ chloramphenicol, and grown overnight at 37°C. Ten mL of the overnight culture were seeded into 1 L LB and grown in the presence of both antibiotics. When the culture reached an OD_600_ ~0.5, 0.8 mM IPTG was added and induction continued at 37°C for 4 hours. Cells were harvested by centrifugation at 3,000 RCF for 10 minutes, and resuspended in 50mL lysis buffer (50 mM Tris-HCl pH 7.6, 150 mM NaCl, 0.5 mM TECP, 10% (v/v) glycerol) with 1x HALT protease inhibitor. Cells were lysed by sonication on ice and centrifuged at 95,000 RCF for 30 minutes. Imidazole was added to the supernatant to a final concentration of 30 mM. A 5 mL HisTrap HP column (GE Healthcare) was pre-equilibrated with 50 mL lysis buffer and the lysate was loaded onto the column and washed with 50 mL of Ni-buffer (50 mM Tris-HCl [pH 7.6], 150 mM NaCl, 10% glycerol (v/v), 30 mM imidazole). PCNA was eluted with a ~110 mL gradient to Ni-buffer + 500 mM imidazole over 120 minutes. PCNλ-containing fractions were identified via 12% SDS-PAGE, dialyzed into storage buffer (20 mM Tris-HCl [pH 7.6], 100 mM NaCl, 10% (v/v) glycerol, 1 mM DTT) and concentrated using a centrifugal filter (10 kDa Amicon, Millipore). Small aliquots were frozen in liquid nitrogen and stored at −80°C.

### Recombineering λ-phage lysogens

Red-based *in vivo* recombination was used to construct recombinant λ-phage DNA (24, 25). First, a lysogen was created by packaging X-phage DNA (cI857ind 1 *Sam* 7; NEB) into empty phage particles (MaxPlax Lambda Packaging Kit, EpiCentre). The packaged phage was used to infect *E. coli* strain 7723 and lysogens were identified via colony PCR and sensitivity to growth at a restrictive temperature (42-45°C). The resulting cells (IF189) were transformed with pKD78, which harbors the Red recombineering system under the control of an arabinose-inducible promoter (26). The insertion cassettes were PCR amplified from helper plasmids (Figure 1) with custom primers (IDT, Table S1) using *Taq* DNA polymerase (NEB# M0320S). PCR reactions were treated with 1 U DpnI (NEB# R0176s) at 37°C for 1 hr and purified by gel extraction (Qiagen Kit #28704) to remove residual plasmid template, which can create false positives during recombineering (25). The gel-extracted DNA was re-suspended in Milli-Q water to a final concentration of 100-150 μg μl^−1^ and used as the targeting DNA for the recombineering reaction.

**Figure 1.**
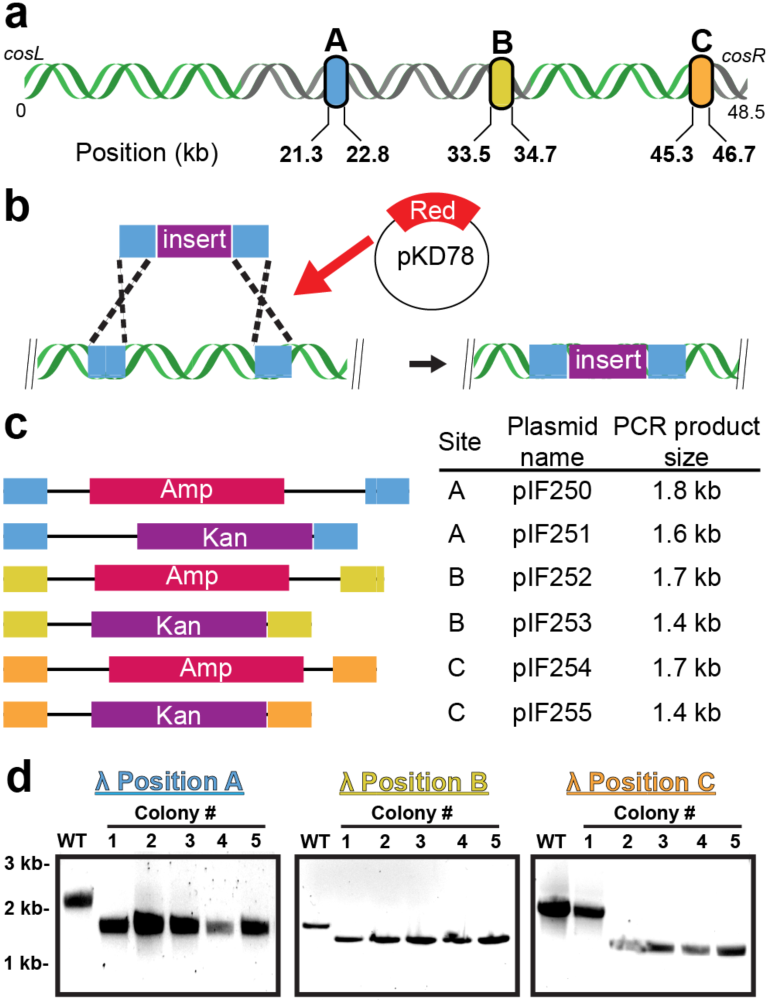
Recombineering at three unique positions within the λ-phage genome. **(a)** Insertion cassettes were designed for three dispensable sites within non-essential regions of the λ-phage genome (shown in gray). These cassettes are 21.3 kb, 33.5 kb and 45.3 kb away from *cosL* (designated A, B and C, respectively). **(b)** Schematic of the Red-based recombineering. An arabinose-inducible plasmid supplies the three Red genes. **(c)** Modules for use as PCR templates to generate cassettes for efficient recombineering. The modules carry an antibiotic resistance gene that is flanked by ~200 bp of homology to each of the sites in (a). **(d)** Recombineering efficiency was scored by colony PCR followed by agarose gels analysis. Successful cassette insertion generates a smaller PCR fragment. We observed 90-100% insertion efficiency at each of the three selected target sites.

Fresh electrocompetent cells were prepared for every recombineering reaction. A 5 mL LB culture of strain IF189 (λ-lysogen) transformed with pKD78 was grown overnight at 30°C in the presence of 10 μg μl^−1^ chloramphenicol. The following day, 350 μl of cells were used to inoculate a fresh 35 mL culture of LB containing the same concentration of antibiotic. When the cells reached an O.D._600_ ~0.5, the Red recombinase system was induced by adding 2% L-arabinose (GoldBio) and incubated for an additional 1 hour at 30°C. Cells were harvested at 4,500 RCF for 7 min, washed three times in ice-cold Milli-Q H**2**O, and finally resuspended in 200 μl of H_2_O (27). Electrocompetent cells were kept on ice and used immediately for the recombineering reaction.

For recombineering, 50-150 ng of targeting DNA was electroporated at 18 kV cm^−1^ in 0.1 cm cuvettes using a micropulser (Biorad #165-210). Cells were immediately resuspended in 1 mL of SOC and then transferred to culture tubes containing 10 mL LB broth. After a 4 hour outgrowth at 30°C, 100 μl of the culture was plated onto LB agar plates containing a low concentration of the appropriate antibiotic. Colonies were checked for successful incorporation of recombinant DNA via colony PCR.

### Purifying phage DNA from lysogens

Expression of recombinant phage was induced via heat shock. A single lysogenic colony was grown in 50 mL of LB broth with the appropriate antibiotic overnight at 30°C. Five mL of this starter culture was used to inoculate 500 mL of LB the following morning. When the flask reached an O.D._600_ ~0.6, the temperature was rapidly raised to 42°C in a water bath. The culture was placed at 45°C in a shaking incubator for 15 minutes and then transferred to a 37°C incubator for two hours. To liberate the phage particles, cells were harvested by centrifugation at 3,000 RCF for 30 minutes and lysed via re-suspension in 10 mL of SM buffer (50 mM Tris-HCl [pH 7.5], 100 mM NaCl, 8 mM MgSO_4_) + 2% chloroform and stirred at 37°C for 30 min at 200 rpm. A subsequent one-hour incubation with 50 ngμl^−1^ DNaseI (Sigma# D2821) and 30 ngμl^−1^ RNaseA (Sigma# R6513) degraded the residual bacterial genomic DNA and RNA. The clarified lysate, containing soluble phage capsids, was obtained by an additional centrifugation for 15 minutes at 6,000 RCF and 4°C, and further diluted with 40 mL of SM buffer. Phage capsids were precipitated during a 1 hour incubation with 10 mL ice-cold buffer L2 (30% PEG 6000, 3M NaCl) and then harvest by centrifugation at 10,000 RCF for 10 minutes at 4°C. The phage pellet was washed with 1 mL of buffer L3 (100 mM Tris-HCl [pH 7.5], 100 mM NaCl, 25 mM EDTA) and then re-suspended with 3 mL of buffer L3, followed by an equal volume of buffer L4 (4% SDS). The phage capsid proteins were further digested by incubation with 100 ngμl^−1^ of proteinase K (NEB # P8012S) for 1 hour at 55°C. SDS was precipitated with 3 mL buffer L5 (3 M potassium acetate [pH 7.5]), and the cloudy solution was clarified by centrifugation at 15,000 RCF for 30 minutes at 4°C. The soluble phage DNA was passed over a pre-equilibrated Qiagen tip-500 column (Qiagen #10262), washed with buffer QC (1.0 M NaCl, 50mM MOPS [pH 7.0], 15% isopropanol) and eluted with 15 mL buffer QF (1.25 M NaCl, 50 mM Tris [pH 8.5], 15% isopropanol). Finally, the DNA was precipitated with the addition of 10.5 mL of 100% isopropanol, rinsed in 70% ethanol and re-dissolved in Te buffer (10 mM Tris-HCl [pH 8.0], 0.1 mM EDTA) to a final DNA concentration of 200-500 ng μl^−1^. We routinely obtained ~250 μg of pure λ-DNA from a single purification.

### Inserting synthetic oligonucleotides into λ-DNA

Recombinant λ-DNA was obtained from strain IF189, which was modified and purified as described above, and 25 μg of the DNA was incubated with 50 U of Nt.BspQI (NEB# R0644s) in a 250 μl reaction with 1X buffer 3.1 (NEB #B7203) at 55°C for 1 hour. The reaction was halted with 1 U of proteinase K (NEB #P8107s) for 1 hour at 55°C. The nicked DNA was mixed with a 100-fold molar excess of the desired insert oligo (e.g., AD005, Table S1), along with a 10-fold excess of *cosL* and cosft-complimentary oligos (IF003, IF004, Table S1). The solution was heated to 70°C for 15 minutes followed by slow cooling to 22°C in a thermocycler at a rate of - 0.5°C min^−1^. The annealed mixture was supplemented with 1000 U of T4 DNA ligase (NEB#M0202L) and 1 mM ATP and further incubated overnight at room temperature in a final reaction volume of 300 μl. From this reaction, 250 μl was further purified by gel filtration while 50 μL was taken for alkaline agarose gel and restriction enzyme digest analysis, as follows.

250 μl of the above reaction was inactivated with high salt (1M NaCl) and then passed through a 120 mL Sephacryl S-1000 column (GE # 17-0476-01), in TE running buffer (10 mM Tris-HCl [pH 8.0], 1 mM EDTA) plus 150mM NaCl, to separate the modified λ DNA from excess oligos and enzymes. The purified DNA was stored at 4°C.

As diagnostics for proper insertion and ligation of our structures into our recombineered λ DNA, agarose gel and restriction enzyme digest analysis was performed, with the proper mock (no insert) and homoduplex (MB032) controls, as described below.

To verify incorporation of the insert, a restriction enzyme digests was performed with NcoI, an enzyme whose recognition site is abolished upon incorporation of the insert into our recombineered λ. 5 λL of the above mentioned ligation reaction was digested with 20 U of NcoI (NEB #R13193) in a 30 μL reaction with 1X Cutsmart buffer (NEB #B7204) for 1 hour at 37°C and run on a 0.8% agarose gel stained with ethidium bromide. To check for complete ligation, a 5 μl aliquot was run over a 0.6% alkaline agarose gel at 15 mV for 20 hours at 4°C. The gel was then submerged in neutralization solution (1 M Tris-HCl [pH 7.6], 1.5 M NaCl) for one hour, and stained in a solution of 10 mg mL^−1^ ethidium bromide for an additional hour. The stained gel was visualized using a Typhoon FLA 9500 laser scanner (GE).

### Imaging Lac repressor on DNA curtains

To obtain distributions of triple FLAG epitope tagged Lac repressor (LacI) binding to recombineered Lac operator λ-DNA, we incubated 120 nM LacI with 10-25 ng μl^−1^ of biotinylated λ-DNA in a 150 [J.L volume of BSA buffer (40mM Tris-HCl [pH 7.6], 50 mM NaCl, 2 mMMgCh, 0.2 mg mL^−1^ BSA, 1 mM DTT) supplemented with 100 mM NaCl, for 5 min on ice. This mixture was diluted to a 1 mL final volume with imaging buffer and injected into flowcells, containing a streptavidin-activated lipid bilayer, to produce single-tethered DNA curtains bound with Lac repressor at the Lac operator site. LacI was then fluorescently labeled *in situ* with an injection of 1.5 nM of anti-FLAG antibodies (Sigma-Aldrich # F9291) conjugated to fluorescent quantum dots (QDs, Life Tech #Q10163MP). The antibodies and QDs were pre-conjugated by mixing in a test tube for 10 minutes on ice prior to injection into the flowcell.

### Loading PCNA on DNA curtains

To observe the binding distribution and diffusion dynamics of PCNA on DNA, 0.8 nM RFC was mixed with 1 nM PCNA (concentrations reported for the trimer) in BSA buffer supplemented with 100 μM ATPγS (Roche). The PCNA/RFC complex was injected into flowcells containing either single‐ or double-tethered DNA curtains, and incubated for 6 minutes at room temperature. Following this incubation, PCNA was labeled *in situ* by injecting 1.5 nM of biotinylated anti-FLAG antibodies conjugated to fluorescent streptavidin-conjugated quantum dots (QDs, emitting at 705 nm, Thermo-Fisher). The antibodies and QDs were pre-conjugated in a test tube on ice prior to injection into the flowcell.

### Single-Molecule Microscopy

Images were collected with a Nikon Ti-E microscope in a prism-TIRF configuration. The inverted microscope setup allowed for the sample to be illuminated by a 488 nm laser light (Coherent) through a quartz prism (Tower Optical Co.). To minimize spatial drift, experiments were conducted on a floating TMC optical table. A 60X water immersion objective lens (1.2 NA, Nikon), two EMCCD cameras (AndoriXon DU897, −80 °C) and NIS-Elements software (Nikon) were used to collect the data with a 200 ms exposure time. Two-color imaging was conducted using a 638 nm dichroic beam splitter (Chroma). Frames were saved as TIFF files without compression for further image analysis in FIJI (NIH).

### Data Analysis

Fluorescent particles were tracked in FIJI with a custom-written particle tracking script (available upon request). The resulting trajectories were analyzed in Matlab (Mathworks). For each frame the fluorescent particle was fit to a two-dimensional Gaussian function to obtain trajectories with sub-pixel resolution (21, 28). To ensure that trajectories corresponded to proteins on DNA, molecules that responded to buffer flow controls were analyzed and DNA-bound proteins were counted for statistical analysis.

## RESULTS

### A molecular toolkit for modifying λ-phage DNA

We sought to develop a method that fulfilled three criteria: (i) multiple exogenous DNA sequences can be inserted into λ-phage DNA with nearly 100% efficiency, (ii) the insertion positions are not limited by availability of unique restriction sites, and (iii) chemical modifications and extra-helical structures can be introduced efficiently and with minimal handling. To satisfy the first design criterion, we surveyed the literature to identify non-essential regions of the λ-phage genome. Two large segments encompassing 37% (17.6 kb) of the genome are dispensable for both lytic and lysogenic growth (Fig. 1a). The first dispensable segment is 16 kilobase (kb) long and is between gene products *J* and *N* (29, 30). The second 1.6 kb segment,is between gene product *Rz* and the *cosR* end (31). Next, we chose three sites—designated sites A, B, and C (Fig. 1a)—because they are each separated by ~12 kb and are thus readily resolved via single-molecule fluorescence microscopy (see below). In sum, these three sites provide convenient site-specific modifications along more than 50% of the length of the entire λ-DNA.

Recombineering is an efficient homologous recombination-based method for genome modification *in vivo* (Fig. 1b) (25, 32). Recombineering proteins, supplied on an arabinose-inducible helper plasmid, catalyze the insertion of a linear DNA substrate between two homologous sites in the phage genome (26). The phage is maintained as an inducible lysogen in *E. coli* (25). We synthesized a series of plasmids with targeting cassettes for sites A, B, and C (Fig. 1c and Table 1). The cassettes consist of a user-specified DNA and an antibiotic resistance gene flanked by ~200 bp of homology to the insertion site. These long homologous regions facilitate efficient recombineering (33). Resistance to the appropriate antibiotic marker further selects the desired product. The recombineering reaction efficiency was scored via colony PCR and agarose gel electrophoresis. Each of the three insertion cassettes replaces a 2.1 kb fragment with a smaller ~1.5 kb insert. As expected, the recombineering efficiency approached nearly 100% at each of the three insertion sites (Fig. 1d, 100%, N=10/10; 100%, N=10/10; and 90%, N=9/10 colonies for sites A, B, and C, respectively). Next, we purified the recombinant λ-DNA by inducing lytic growth at a non-permissive temperature (> 42°C). Above 42°C, unfolding of the temperature-sensitive repressor (*cI857ind 1*) induces phage replication and packaging (30, 34). An additional amber mutation in the S gene (*Sam 7*) delays cell lysis and produces large burst sizes (>200 phage particles per cell), increasing the overall yield of phage particles (29, 35). We purified the recombinant λ-DNA from phage and confirmed site-specific insertion of the specified DNA via PCR and restriction enzyme digestion (Fig. 1). In sum, we developed a series of drug-selectable cassettes for rapid and site-specific insertion of recombinant DNA into the λ-phage genome.

As a first proof-of-principle, we used high-throughput DNA curtains to measure the binding positions of individual Lac repressor (LacI) proteins on recombinant λ-DNA harboring a high-affinity Lac operator sequence in one of the three target sites. In this assay, a fluid lipid bilayer is deposited on the surface of a micro-patterned flowcell (36–38). Biotinylated DNA is immobilized on the lipid bilayer via a biotin-streptavidin linkage and buffer flow is used to organize hundreds of DNA molecules at chromium (Cr) diffusion barriers (Fig. 2a) (36, 38). Epitope-labeled LacI was pre-incubated with the DNA and fluorescently labeled with anti-FLAG antibody-conjugated quantum dots (QDs). The DNA was stained with a fluorescent intercalating dye (YOYO-1) and 89% (N=89/100) of the DNA molecules where full-length (Fig. 2b). This result likely underestimates the overall quality of the recombinant DNA, as both shear forces and YOYO-1 induced DNA damage likely contributing to the observed broken DNA molecules. As expected, Lac repressor bound specifically to the recombinant operator site (Fig. 2c) at each of our three target sites (39). These results demonstrate that high quality recombinant λ-DNA provides an ideal substrate for single-molecule imaging.

**Figure 2.**
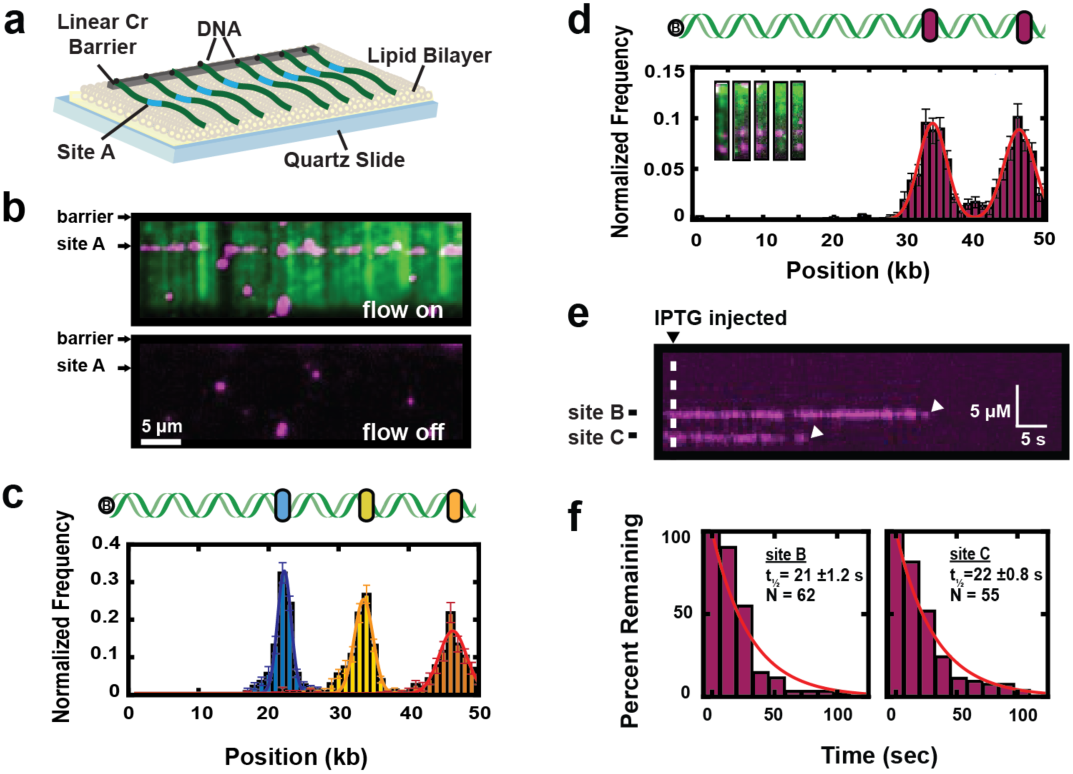
Single-molecule imaging of Lac repressor dynamics on recombineered λ-DNA substrates. **(a)** Schematic of a single-tethered DNA curtain assay. DNA molecules (green) are immobilized on a fluid lipid bilayer and arranged at micro-fabricated Chromium (Cr) barriers. One of the insertion sites (site A) is indicated in blue. **(b)** TIRFM image of Lac repressor (magenta) binding to a lac operator inserted at site A on a recombineered λ-DNA substrate (green). **(c)** Binding distributions of Lac repressor to a lac operator sequence inserted at one of the three sites within λ-DNA. Solid lines indicate a Gaussian fit to the data. Blue: position A (21.7 ± 1.0 kb, N=558, blue). Orange: position B (34.0 ± 1.5 kb, N=538). Yellow: position C (46.0 ± 2.3 kb, N=467). Error indicates the standard deviation of the Gaussian fit. **(d)** Insertion of two lac operator cassettes within a single λ-DNA construct. Inset: single-molecule imaging of the recombinant DNA shows Lac repressor at both positions B & C. Binding histograms show nearly equal occupancy at both lac operator sites. Red line: double Gaussian fit (34.0 ± 2.0 kb and 46.2 ± 2.2 kb, respectively; N = 525). **(e)** As expected, lac repressor dissociates from both operator sites after injection of 0.4 mM Isopropyl β-D-1-thiogalactopyranoside (IPTG). **(f)** Quantification of the Lac repressor lifetimes upon IPTG injection. Solid lines: single-exponential fits (t_½_ = 21.3 ± 1.2 sec, N=62 and 22 ± 0.8 sec, N=55 at positions B & C, respectfully).

Next, we developed a strategy for inserting multiple recombinant DNA sequences—each flanked by a different antibiotic cassette—into the same λ-DNA substrate. A series of targeting vectors were constructed with either an ampicillin or kanamycin resistance gene flanked by ~200bp of homology to sites A, B or C on the phage genome (Fig. 1c). Using these cassettes, we tested the efficiency of incorporating a Lac operator into both sites B & C of the same DNA substrate (Fig. 2d). Recombineering efficiency was scored by colony PCR, with 100% (N=10/10) of all colonies harboring both insertion cassettes. To confirm the quality of these DNA substrates for single-molecule experiments, we imaged Lac repressor binding to DNA curtains harboring a Lac operator in both sites B & C. As expected, 92% (N=483/525) of all Lac repressor molecules bound to either one of the two insertion sites (Fig. 2d). Furthermore, injecting 1 mM IPTG into the flowcell rapidly dissociated all LacI molecules with a half-life of 21±1.2 seconds (N=65; error indicates 95% C.I.) for site B and 22±0.8 seconds (N=55) for site C (Figs. 2e & 2f).

The broad LacI binding distributions largely stems from experimental uncertainty in the precise surface tethering of DNA molecules (Figs. 2c & 2d). To construct binding histograms, we assume that all DNA molecules are tethered at the center of the diffusion barriers, which are ~ 1-1.5 μm wide (36). When compiling binding distributions from hundreds of such DNA molecules (spanning several flowcells), this approximation results in an apparent broadening of the resulting histogram. This broadening has also been observed for both site‐ and sequence-specific DNA-binding proteins in similar flow-stretches assays (e.g., EcoRI(E111Q) (40, 41), FtsK (42, 43), and DNA repair proteins (44, 45)). Nonetheless, these results confirm that high-quality, recombinant λ-DNA can be rapidly modified at multiple sites and purified for single-molecule studies.

### Insertion of modified oligonucleotides into λ-DNA

Single-molecule studies frequently require the insertion of modified bases or fluorophores at precise positions along the DNA substrates. To meet this need, we developed an improved nicking-enzyme based strategy for inserting synthetic oligonucleotides at precise positions along λ-DNA (Fig. 3a). We designed a nicking cassette that includes three consecutive BspQI recognition sites separated by 20 bp spacers. The DNA is nicked with Nt.BspQI (nickase) enzyme that creates closely spaced discontinuities along one of the two DNA strands. λ-DNA harbors ten widely spaced BspQI recognition sites, thereby limiting nicks and double-strand breaks along the rest of the DNA substrate. The nicked DNA is heated to liberate short single-stranded DNA fragments. A synthetic oligonucleotide is inserted into the backbone by annealing and ligation (9). After nicking with Nt.BspQI and heating, the gapped DNA is slowly cooled with a 100-fold excess of the desired insertion oligo. The annealed and ligated λ-DNA is further purified by gel filtration to remove enzymes and excess oligonucleotides. Conversion of restriction sites (NcoI, NotI) to single-stranded DNA within the nicking cassette allow rapid quantification of the insertion efficiency via restriction digest mapping.

**Figure 3.**
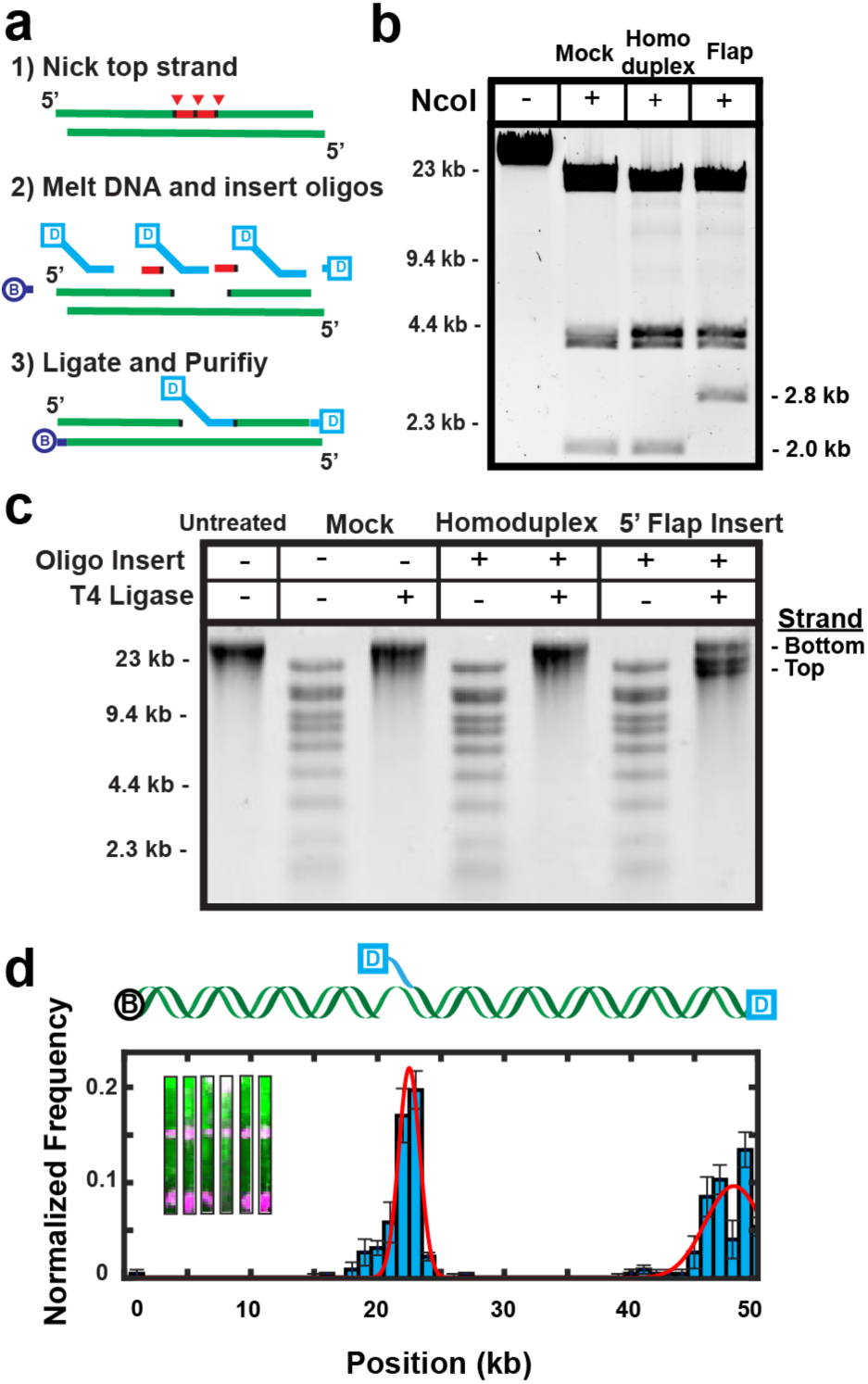
Construction of DNA substrates containing extrahelical structures. **(a)** Schematic of nickase-mediated insertion of a synthetic oligonucleotide (Digoxigenin (dig)-labeled 5’-flap). **(b)** Diagnostic digest of a recombineered λ, demonstrated here by the 2.7 kb band in lane 4 which is digested into two smaller bands in lanes 2 and 3 (bottom band not shown). **(c)** Denaturing alkaline agarose gel of three separate reactions (no insert, mock insert, flap insert) to determine ligation efficiency. The last lane shows a larger band (bottom strand) and two smaller bands (top strand) that are created upon insertion of the flap. **(d)** Distribution of quantum-dot labeling of the 5’ flap (magenta, 22 ± 0.8 kb) and the Dig-modified 3’ end (magenta, 47.5 ± 2 kb) of the λ DNA construct (green), N=225. The insets show single DNA (green) labeled with quantum dots (magenta) at position A and at the free DNA ends.

We validated this approach by inserting extrahelical structures into a user-defined site (site A) within λ-DNA. The nicking cassette was cloned into λ-DNA using the recombineering strategy described above. Next, we inserted a 5’-ssDNA flap oligonucleotide that annealed to 26 bp of the gapped ssDNA. The incorporation efficiency was scored via cleavage with NcoI (Fig. 3b). Insertion of the flap oligonucleotide abolishes a single NcoI site within the cassette, converting two small NcoI DNA fragments to a larger 2.8 kb band on a native agarose gel (Fig. 3b). As expected, we saw 100% cleavage of the flap-proximal NcoI site with both untreated and mock-treated DNA substrates, whereas ~1% of the DNA molecules were Ncol-sensitive after flap incorporation. Next, alkaline agarose gel electrophoresis was used to assay the quality of the ligated DNA substrate (Fig. 3c). The 48.5-kb DNA runs as a single large band on a denaturing gel. Nicking with Nt.BspQI produces a distinct ladder of bands, which are removed after ligation. As expected, the resulting DNA substrate shows three distinct bands on the alkaline agarose gel; the full-length bottom strand and the 21.3 kb and 26.7 kb discontinuous top strands. Importantly, ligating the nicked DNA substrate with a complimentary phosphorylated oligonucleotide produces a substrate that is indistinguishable from the unmodified λ-DNA (Fig. 3c). Together, these ensemble-biochemical results confirm the efficient and site-specific incorporation of a synthetic oligonucleotide into the nicking cassette in λ-DNA.

In addition to the biochemical assays, we also confirmed the overall quality of the DNA substrate and site-specific flap incorporation via single-molecule fluorescence imaging (Fig. 3d). We assembled single-tethered DNA curtains with λ-DNA that has a biotin on the *cosL* end and a digoxigenin on both the inserted flap and the *cosR* end. Digoxigenin was fluorescently labeled via anti-digoxigenin-functionalized QDs. The fluorescent signal was localized to the flap and DNA end regions (Fig. 3d, inset). A histogram of the binding positions revealed two peaks, one at the distal DNA end and a second at site A, where the ssDNA flap was incorporated (Fig. 3d). Importantly, 82% (N=82/100) of the fluorescently stained DNA molecules were full-length, confirming that the DNA was not fragmented after the oligo insertion reaction. In sum, our ensemble biochemical and single-molecule results confirm efficient and site-specific incorporation of synthetic oligonucleotides into full-length λ-DNA.

### Observing PCNA dynamics on modified DNA substrates

To further demonstrate the utility of modified λ-DNA substrates for single-molecule studies, we imaged the specificity and diffusion dynamics of *S. cerevisiae* proliferating cell nuclear antigen (PCNA) on recombinant DNA curtains. For fluorescent labeling, PCNA was over-expressed with a triple FLAG epitope tag on the N-terminus, which is far away from the DNA binding site (46, 47). PCNA is loaded onto DNA by the replication factor C (RFC) complex. RFC uses the energy of ATP hydrolysis to open and close the PCNA sliding clamp onto DNA during synthesis and repair (46, 47). Therefore, we also purified recombinant *S. cerevisiae* RFC (22). After incubating and co-injecting the RFC-PCNA complexes into the flowcell, PCNA was fluorescently labeled *in situ* with anti-FLAG functionalized quantum dots (QDs), as described previously for the human RFC-PCNA complex (6). We titrated down the concentration of both RFC and PCNA to load one or fewer PCNA complexes per flow-stretched λ-DNA molecule. As expected the loading reaction was both RFC‐ and ATP-dependent. Following PCNA loading, ATP hydrolysis catalyzes the release of RFC from DNA. However, to ensure complete RFC removal, we stringently washed the flowcells with buffer containing 300 mM NaCl. Following RFC removal, we observed freely diffusing PCNA molecules on the double-tethered DNA curtains (Fig. S1). QDs increase the effective PCNA cargo (~10 nm radius), but the fluorescent PCNA complexes retain full diffusive activities on homoduplex DNA complex (Fig. S1 and ref. (6)). As demonstrated previously for the human PCNA complex, the diffusion coefficient of QD-labeled yeast PCNA is only moderately dependent on the ionic strength and is in quantitative agreement with the values obtained for the human protein (Fig. S1C). These observations demonstrate both loading and fluorescent tracking of individual yeast PCNA complexes on homoduplex DNA.

Next, we explored the PCNA loading specificity by full-length *S. cerevisiae* replication factor C (RFC) clamp loader. Full-length RFC catalyzes nonspecific PCNA loading on DNA, and this activity is stimulated by a homoduplex DNA-binding motif within the large subunit (Rfc1) of RFC (48–50). Due to this non-specific loading activity, most previous studies have used a truncated RFCΔN, where the ATP-independent DNA-binding patch of Rfc1 has been deleted (51). More recently, RFCΔN has also been proposed to directly load PCNA on extrahelical triplet nucleotides repeats (TNRs), ultimately driving TNR expansion (19, 20). However, the loading specificity of PCNA by the full-length RFC complex has not been directly observed. Thus, we developed an assay that can resolve the relative loading efficiencies of PCNA by RFC on both homoduplex and extrahelical DNA.

We analyzed PCNA loading by RFC on three modified DNA substrates harboring a 30-nt 5’-ssDNA flap which resembles a primer-template junction, a 13xCAG repeat, or a mock-treated homoduplex DNA (Fig. 4). To capture the RFC-PCNA complex at the loading site, ATP was substituted for ATPγS, a slowly-hydrolyzable ATP analog. ATPγS traps the RFC-PCNA complex at the loading site and prevents PCNA release and ring closing (46, 51, 52). As expected, RFC preferentially loads PCNA at the 5’-ssDNA flap (inserted into site A), which resembles a primed DNA site (Figs. 4a & 4b). RFC also nonspecifically loads PCNA on the 48.5 kb homoduplex DNA substrate, which is consistent with prior studies (51-53). Turning off buffer flow retracted both PCNA and the associated DNA molecules to the top barrier, confirming that PCNA was loaded on DNA (Fig. 4a, bottom). We also prepared a recombinant DNA substrate with a 13xCAG repeat or a homoduplex oligo (mock insertion) instead of the 5’-ssDNA flap at site A. We observed modest enrichment of PCNA on a 13xCAG repeat (Fig. 4b, middle) relative to mock-treated homoduplex DNA (Fig. 4b, bottom). To better characterize the loading efficiency on these structures, we calculated the relative enrichment of PCNA in a 5 kb region spanning the nicking cassette on the three recombinant DNA molecules (Fig. 5d). This region was selected because it is expected to capture ~99% (~3 standard deviations) of all site-specifically bound PCNA molecules (within our spatial resolution). As expected, the flap structure showed 2.7-fold enrichment over homoduplex DNA, whereas loading at the 13xCAG was 1.5-fold higher than mock-treated homoduplex DNA. These results are consistent with recent ensemble biochemical studies that inferred that a truncated *S. cerevisiae* RFCΔN can load *H. sapiens* PCNA on 3xCAG repeats (19). Therefore, these results demonstrate that full-length RFC preferentially loads PCNA on flap and 13xCAG extrahelical structures relative to homoduplex DNA.

**Figure 4.**
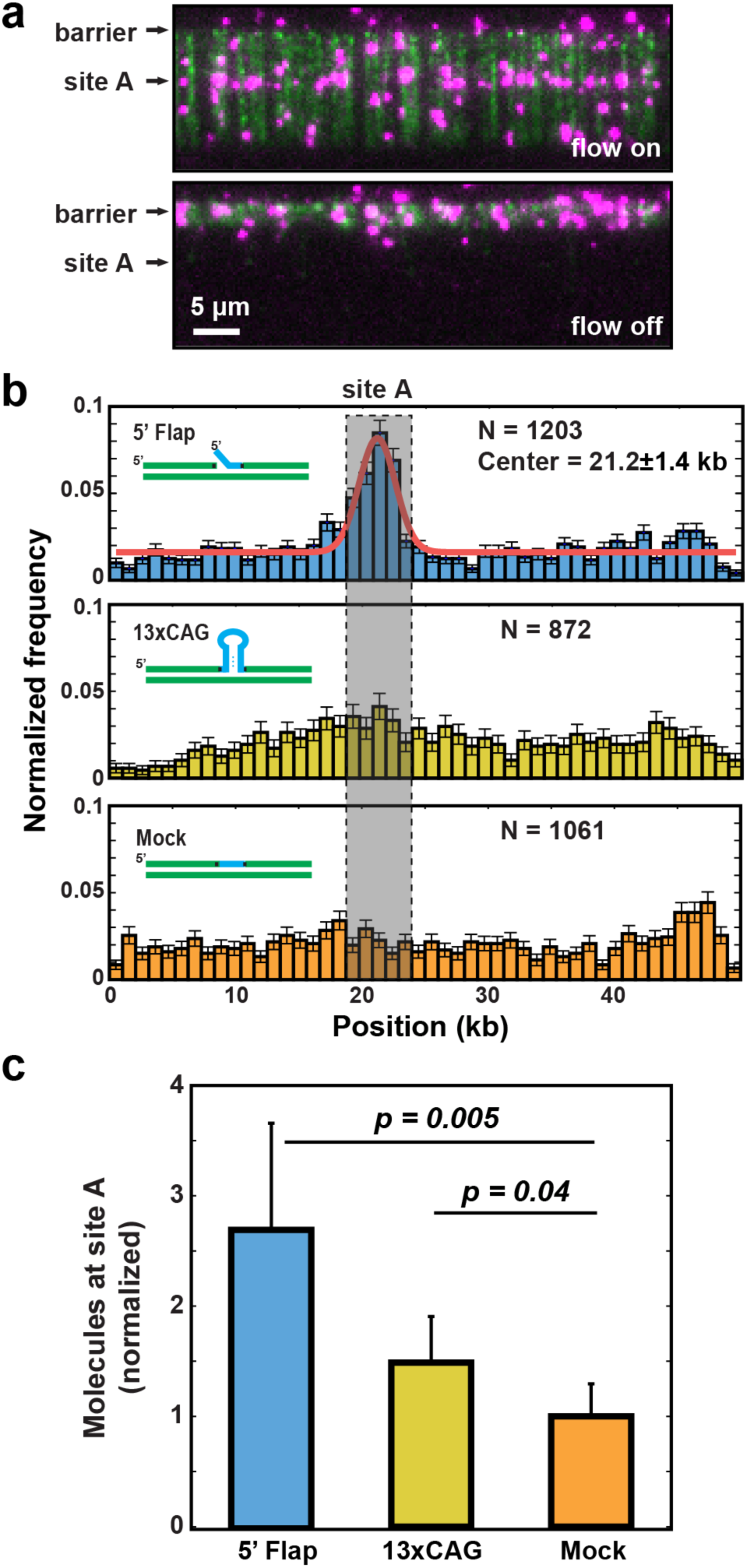
RFC loads PCNA on various extrahelical DNA structures. **(a)** Fluorescent images of PCNA on DNA substrates with a 5’-ssDNA flap inserted at site A. Top: with buffer flow, bottom: without flow. DNA is stained with YOYO-1 (green) and PCNA is labeled with anti-FLAG antibody conjugated QDs (magenta). Terminating buffer flow retracts both PCNA and DNA to the Cr barrier, confirming that PCNA is bound to the DNA and is not on the surface (bottom panel). (b) Binding distribution histogram of PCNA/RFC on λ-DNA molecules containing a 5’-ssDNA flap (top), 13xCAG repeat (middle), and mock-treated homo-duplex DNA (bottom). The binding distribution on a 5’-ssDNA flap-containing DNA was fit to a Gaussian distribution (red line, top panel). The Gaussian fitting shows the center of 21.2 kb ± 1.4 kb (st. dev.), confirming that RFC prefers to load PCNA on ss/dsDNA junctions. Gray box: a 5 kb-wide region, as determined by the center and ~3 standard deviations of the Gaussian fit of the 5’-ssDNA flap structure. (c) The mean number of PCNA molecules loaded at site A (within the gray box), as determined by bootstrap analysis of the histograms in (b). Error bars represent the st. dev. of the mean from the bootstrap analysis. There was a statistically significant enrichment of PCNA at both 5’-ssDNA flaps and 13xCAG repeats relative to mock-treated homoduplex DNA.

**Figure 5.**
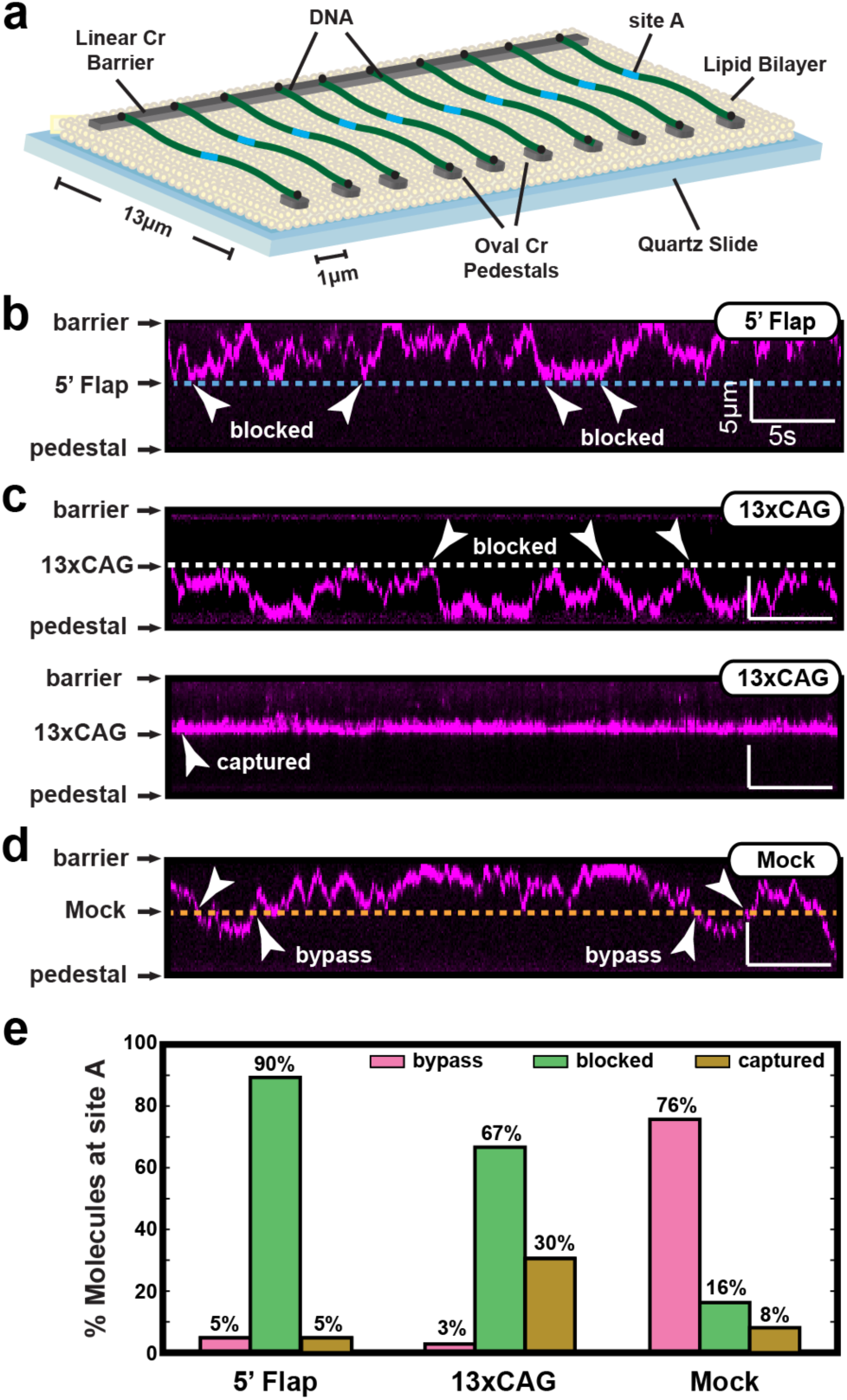
PCNA diffusion on λ-DNA containing various extrahelical structures. (a) An illustration of the double-tethered DNA curtains. Oval chromium (Cr) pedestals are coated with anti-digoxigenin antibodies, and buffer flow is used to immobilize the dig-labeled λ-DNA between the linear barriers and Cr pedestals. Kymograph of diffusing PCNA molecule on DNA substrates with (b) a 5’-ssDNA flap, (c) a 13xCAG repeat, or (d) mock-treated homoduplex DNA. The characteristic changes in diffusion behavior or PCNA are indicated with arrows, and the dashed lines indicate site A. (e) Percentage of molecules showing either bypass, blocked, or captured behavior at site A. At least 25 DNA molecules were analyzed and classified into each of three categories (N=40, 36, and 37 for the flap, 13xCAG, and mock DNA substrates).

Next, we observed PCNA diffusion in the context of extrahelical DNA structures. PCNA has a 3.4 nm-diameter inner opening that is larger than the width of B-form DNA (46, 48, 49). The large inner opening of the PCNA trimer facilitates sliding past small mismatches and bulges, but may be blocked by larger extrahelical structures (50). To directly test this proposal, we used double-tethered DNA curtains to observe PCNA diffusion on various DNA substrates (Fig. 5a). In this assay, one end of the DNA was labeled with a biotin and the second end was labeled with a digoxigenin (dig). Pedestals that were located 13 μm away from the diffusion barriers were decorated with anti-dig antibodies. The antibody-coated pedestals provide a second attachment point for dig-labeled DNA molecules (36, 38). The DNA molecules that are tethered between the barriers and pedestal remain fully extended without the need for additional buffer flow (Fig. 5a). Using double-tethered DNA molecules, we monitored PCNA diffusion on DNA substrates containing a 5’-ssDNA flap, a 13xCAG, as well as mock-treated homoduplex DNA (Fig. 5b-d).

We first observed PCNA diffusion dynamics on DNA harboring a 5’-ssDNA flap (Fig. 5b). PCNA can approach the flap from one of two orientations: (i) in the *cosL* ➔ *cosR* direction PCNA encounters a 4-nt ssDNA gap prior to the flap and (ii) in the *cosR* ➔ *cosL* direction PCNA encounters the 5’-T30 tail. Out of 40 PCNA molecules 90% (N=36/40) were blocked by the flap in either orientation (N=18 for *cosL* ➔ *cosR* and N=18 for *cosR* ➔ *cosL*). The remaining molecules were either captured at the nickase cassette (5%, N=2/40) or directly bypassed (5%, N=2/40) the flap structure. This observation may stem from incomplete re-ligation of the oligo or from other DNA lesions that may exist in the 48kb-long DNA structure. Nonetheless, these data suggest that that the flap is too large to be accommodated within the PCNA pore.

Next, we studied PCNA diffusion on DNA harboring a 13xCAG repeat (Fig. 5c). As with the flap, PCNA that loaded on homoduplex DNA could not diffuse across the 13xCAG repeat (97%, N=35/36). Remarkably, we also observed that 30% (N=11/36) of the PCNA molecules that loaded directly at the 13xCAG structure were completely stationary, without exhibiting any diffusive properties. This result is consistent with a mechanism where RFC loads PCNA within the 13xCAG repeat, trapping the clamp at that lesion (19). Finally, mock-treated DNA supported PCNA diffusion over the complete length of the DNA molecule. Diffusion molecules were largely unimpeded by oligo replacement within the nicking cassette (Fig. 5d). We also observed a few stationary PCNA molecules (N=3/37, 8%) at the mock-treated nicking cassette. In addition, 16% (6/37) of the diffusing PCNA molecules could not immediately traverse the mock-treated site. These observations may stem from incomplete re-ligation of the homoduplex oligo or from a small ssDNA gap that may occur if the annealing was incomplete. Nonetheless, the large differences in the PCNA diffusion on the 5’-ssDNA flap, the 13xCAG repeat, and the mock-treated DNA confirm that PCNA diffusion is blocked by large extrahelical structures and that 13xCAG repeats can capture PCNA, consistent with a role for trapped PCNA in promoting TNR expansion. We conclude that the nickase-based approach developed here permits the insertion of synthetic oligonucleotides at defined positions within λ-DNA.

## DISCUSSION

The results show that a recombineering-based molecular toolbox facilitates rapid and site-specific incorporation of exogenous DNA sequences and structures in multiple positions in λ-DNA. As a proof-of-principle, we showed site-specific insertion of a 1-kb cassette at three positions along the λ-DNA. Both larger and smaller modifications of the λ-DNA molecule are also possible, provided that nonessential regions of the phage genome are targeted for modification (Fig. 1). For such applications, we developed a series of ampicillin‐ and kanamycin-resistant cassettes that can accommodate large insertion fragments that are flanked by regions of homology to three distinct locations within the phage DNA. These markers facilitate positive selection, ultimately yielding nearly 100% recombination efficiencies. If required, the drug markers can be removed by flanking the cassettes with FRT sites and inducing Flprecombinase*in vivo* (25, 26). Here, we focused on a series of recombinant λ-DNAs that maintained near-wild type length. However, the DNA substrate may be shortened or expanded, provided that the λ-DNA remains within 78%-105% (38 to 53 kb) the length of the wild-type phage genome. This requirement is essential for efficient packaging of the DNA into phage capsids (30). This flexibility guarantees that both large and small insertions can be made along the non-essential phage genome.

We regularly achieved 90-100% recombineering efficiency using PCR constructs directed towards one of three unique sites within the λ-phage genome. However, we also noticed that deviating from a few best practices could produce low yields or off-target insertion products. We summarize these observations below. First, cryopreserving lysogens with the recombineering helper plasmid resulted in high off-target recombination rates. Substantially higher yields where achieved when the lysogens were prepared fresh and used immediately. Secondly, we induced the Red system in mid-log cells (at an OD_600_~0.5 or below), as the insertion efficiency was greatly reduced when cells approach stationary phase. Additionally, recombineering efficiency was higher when the PCR products were DpnI-treated to remove residual template DNA, gel purified, and prepared at a concentration of 100 ngλl^−1^ or greater. After electroporation, a minimum 4-hour outgrowth should be allowed before plating the cells on antibiotic selective plates. Finally, recombineering at site A yielded slow-growing cells that required an overnight outgrowth in LB and up to two days for colonies to appear on antibiotic selective plates. Following these simple guidelines routinely resulted in highly specific modification at the desired locus.

We also demonstrated an improved nickase-based strategy for incorporating extrahelical structures at defined positions along the DNA duplex. Nickase-based DNA modifications require designer nicking cassettes with closely-spaced nicking sites (9). Although this approach is most commonly used with plasmid-based modifications, it has also been demonstrated for modifying a single locus within λ-DNA (9, 15, 18). However, the requirement for two or more closely spaced nicks required a nicking enzyme that cleaves λ-DNA at hundreds of sites on both strands, leading to double-stranded breaks and low yields of full-length DNA (15, 18). Our approach refines the previously published strategies in three key ways: (i) the nicking cassette can be inserted in any non-essential region of the genome, (ii) nicking at three adjacent sites exposes 39 nt of single stranded DNA, providing sufficient space to insert large oligonucleotides (e.g., long ssDNA gaps, RNA-DNA structures, and other chemically modified oligonucleotides), (iii) using a nickase with few recognition sites minimizes damage to the DNA substrate, and (iv) incorporation efficiency can be rapidly scored via restriction enzyme digestion with one of several enzymes. Finally, the 39-nt gap can be replaced by a single long synthetic oligo, which is thermodynamically more stable than the multiple short fragments that are removed after nicking. Together, these considerations maximize the efficiency of the replacement reaction.

Using this approach we demonstrated that *wt* RFC preferentially loads PCNA on 5’-ssDNA flaps and 13xCAG repeats relative to homoduplex DNA. PCNA can diffuse on DNA but is blocked by both flaps and 13xCAG repeats, consistent with the 3.4 nm radius of the PCNA pore (50). Finally, PCNA can be directly loaded onto the 13xCAG repeat, preventing further movement away from the lesion. These results shed light on PCNA diffusion dynamics and further highlight the utility of this molecular toolkit for both single-molecule and ensemble biochemical studies.

## ACKNOWLEDGMENTS

We are indebted to Manju Hingorani, Bruce Stillman, and Francisco Blanco for sharing recombinant plasmids. We are grateful to members of the Finkelstein lab for carefully reading the manuscript. The content is solely the responsibility of the authors and does not necessarily represent the official views of the National Institutes of Health.

